# *In Vitro* and *In Vivo* activity of a novel catheter lock solution against bacterial and fungal biofilms

**DOI:** 10.1101/300145

**Authors:** J Chandra, L Long, N Isham, PK Mukherjee, G DiSciullo, K Appelt, MA Ghannoum

## Abstract

Central line associated bloodstream infections (CLABSIs) are increasingly recognized to be associated with intralumenal microbial biofilms, and effective measures for the prevention and treatment of BSI remain lacking. This report evaluates a new commercially developed antimicrobial catheter lock solution (ACL) containing trimethoprim (5 mg/ml) and ethanol (25%) and CA-EDTA 3% for activity against bacterial and fungal biofilms using *in vitro* and *in vivo* (rabbit) catheter biofilm models. Biofilms were formed with bacterial (seven different species including vancomycin-resistant enterococcus, VRE) or fungal (*C*. *albicans*) species on catheter materials. Biofilm formation was evaluated by quantitative culture (colony forming units, CFUs) and scanning electron microscopy (SEM). Treatment with ACL inhibited growth of adhesion phase biofilms *in vitro* after 60 min (VRE) or 15 min (all others), while mature biofilms were eradicated after exposure for 2 or 4 h, compared to control. Similar results were observed for drug-resistant bacteria. In the catheterized rabbit model, when compared against heparinized saline control, ACL lock therapy significantly reduced the catheter bacterial (3.49 ± 0.75 vs. 0.03 ± 0.06 log CFU/catheter, respectively; *P* = 0.001) and fungal burden (2.48 ± 1.60 vs. 0.55 ± 1.19 log CFU/catheter segment, respectively; *P* = 0.012). SEM also demonstrated eradication of bacterial and fungal biofilms *in vivo* on catheters exposed to ACL, while vigorous biofilms were observed on untreated control catheters. Our results demonstrate that ACL was efficacious against both adhesion phase and mature biofilms formed by bacteria and fungi *in vitro* as well as *in vivo*.

## INTRODUCTION

Central venous catheters (CVCs) are intravascular devices inserted centrally or peripherally and terminate at the heart or in a proximal great vessel. CVCs are indispensable tools in the acute hospital care and outpatient settings that enable infusion therapy such as fluid management, i.v. administration of drugs, cancer chemotherapeutics, hemodynamic monitoring, parenteral nutrition and hemodialysis. CVCs enable continuous access to the central venous system without the need for repeated venipuncture (1).

All intravascular catheters communicate with the vascular system and therefore must be filled with a physiologically compatible solution to create a hydrostatic lock between the catheter hub and the circulatory system. Lock solutions physically oppose the retrograde flow of blood back into the indwelled catheter, however in doing so they support microbial growth and biofilm formation inside catheters that are isolated from immune surveillance. Biofilms are complex three-dimensional structures composed of microorganisms living in a microbial generated extracellular matrix of proteins, nucleic acids, and polysaccharides, and are characteristically less-susceptible to antimicrobial agents (2, 3). Biofilms formed by bacteria (e.g. *Staphylococcus aureus*, *S*. *epidermidis*, *Pseudomonas aeruginosa* and *Acinetobacter* spp.) as well as fungi (e.g. *Candida albicans*) have the potential to go through cycles of active shedding, thus creating a nidus of systemic pathogen dissemination that may be the cause of recurring blood stream infections and sepsis (2 – 9).

Worldwide the standard of care in lock solutions is phosphate buffered saline (PBS) or PBS containing heparin. Heparin is a polysaccharide that is used as an anticoagulant to prevent blood coagulation within the catheter lumen (10). Although Heparin is a potent anticoagulant, it is associated with blood stream infections and bleeding complications (11).

Chronically and critically ill patients with indwelled CVCs are at risk of developing CLABSI, which contribute to patient morbidity, mortality, extended length of hospital stay, and increased cost of care (12 – 17). According to the Agency for Healthcare Research and Quality in the U.S. Department of Health and Human services, CLABSIs are responsible for an estimated 84,551 to 203,916 preventable infections at a cost of $1.7 to $21.4 billion annually (18). Mortality rates from diagnosed CLABSIs range from 12-25%, according to the National Healthcare Safety Network (19).

In recent years, several antimicrobial catheter lock solutions have been described and clinically evaluated. Meta analyses of human clinical trials involving off-label compounded antibiotic lock solutions have demonstrated statistically significant reductions in catheter blood stream infections (20), as has taurolidine containing lock solutions (21).

In this report we describe a new commercially prepared antimicrobial catheter lock solution containing the antibiotic trimethoprim 5 mg/ml and ethanol 25% v and CA-EDTA 3% in a physiologically balanced phosphate buffered saline solution (22, 23).

This study evaluated the activity of ACL against early adhesion phase and 24-hour mature phase bacterial and fungal biofilms using *in vitro* and *in vivo* (rabbit) catheter biofilm models. The data demonstrate that ACL is efficacious against both adhesion phase and mature biofilms formed by bacteria and fungi.

## MATERIALS AND METHODS

### Organisms and Reagents

Organisms tested for the catheter-associated biofilm model were *Escherichia coli*, *Enterococcus faecium*, *Enterobacter cloacae*, *P*. *aeruginosa*, methicillin-resistant *S*. *aureus* (MRSA), vancomycin-resistant enterococcus (VRE), *Serratia marcescens* and *C*. *albicans* taken from the strain collection of the Center for Medical Mycology. The antimicrobial catheter lock solution containing trimethoprim 5 mg/ml, ethanol 25% v/v, and 3% calcium EDTA was prepared according to good manufacturing practices (GMP) by AAI Pharma Services Corporation, now Alcami, Inc.

### Biofilm formation

Early adhesion and mature phase biofilms were formed according to our standardized protocol described previously (2, 4). Fungal and bacterial biofilms were grown on silicone elastomer (SE) disks with a 1.5 cm diameter (Cardiovascular Instrument Corp., Wakefield, Mass.). To facilitate biofilm growth and adhesion, sterile disks were placed into 12 well tissue culture plates containing 2 mls of fetal bovine serum (FBS) each. Plates were placed on a rocker and incubated for 24 h at 37°C. Disks were then removed and washed with phosphate-buffered saline (PBS) to eliminate any remaining FBS. For the formation of early (90 min) adhesion phase biofilms, FBS coated disks were then soaked in 3 ml of 1 × 10^7^ cells/ml and incubated for 90 minutes at 37°C. After 90 minutes, wells were gently washed with PBS to remove nonadherent cells. Following adhesion phase biofilm formation, discs were incubated for 15, 30 or 60 min in the presence of ACL (4 ml) or PBS control.

For the evaluation of activity against mature biofilms, bacterial and fungal cultures were allowed to establish mature biofilms for an additional 24 h (bacteria) or 48 h (*Candida*). Mature biofilms of bacteria and *Candida* were then exposed to 4 ml of phosphate buffered saline (PBS) control or ACL for 2 and 4 h.

### Activity of lock solution against microbial biofilms *in vitro*

At each time point, aliquots of adhesion and mature phase PBS treated control biofilms, as well as and those exposed to ACL were harvested. SE discs were then scraped and the scraped material was suspended in 100 μl. Ten-fold serial dilutions were made and spread on PDA or BHI agar plates to enable microbial growth and obtain CFUs. Plates were incubated for 24 h at 37°C and CFUs counted. Discs incubated with PBS alone, were used as controls.

### Efficacy of catheter lock solution *in vivo*

ACL was also tested against biofilms formed by *C*. *albicans* or MRSA *in vivo* using our previously described rabbit lock therapy model (24). A standard inoculum (300 μl) consisting of 10^7^ CFU/ml of *C*. *albicans* (M61) or MRSA (ATCC 43300) was prepared in sterile normal saline with 100 U of heparin (Abbott Laboratories, North Chicago, IL).

Female New Zealand white rabbits weighing between 2.5 to 3.5 kg (Covance, Inc., Princeton, NJ) were housed at the Case Western Reserve University Animal Facility, adhering to Institutional Animal Care and Use Committee guidelines. After the animals were allowed to acclimatize for 7 days, surgery was performed to place a silicone catheter in the external jugular vein. The catheters were made with silastic tubing cut to 30 cm, with an internal diameter of 0.04-in. and an external diameter of 0.085- in. (Dow Corning, Midland, Mich.). To protect the catheter from collapse and ensure that it was properly positioned, a polyethylene cuff (PE 240; Becton Dickinson, Sparks, Md.) was slipped over the catheter and superglued 4 cm from the inserted end. Rabbits were anesthetized intramuscularly using a cocktail of ketamine and xylazine prior to surgery. The animals were clipped, and a surgical scrub was performed. Silicone catheters were placed in the right external jugular vein and tunneled subcutaneously as described previously (24). Catheters were inoculated with 10^7^ CFU/ml of *C*. *albicans* or MRSA, which was ‘locked’ in the lumen of the catheter for 24 h to allow the formation of intralumenal biofilm. Free floating organisms were then by aspirating the inoculum broth and daily heparinized saline flushes, 100 U of heparin in sterile normal saline, were performed for 3 days. Once mature biofilms were formed, blood samples were obtained from the catheter and submitted for culture to confirm the presence of MRSA or *C. albicans*. Animals inoculated with MRSA were randomized into the following groups: 1) ACL lock therapy for 2 h/day for 7 days (n = 4), and 2) untreated control (n = 5). Similarly, animals inoculated with *C*. *albicans* were randomized into the following groups: 1) ACL lock therapy for 2 h/day for 7 days (n = 16), and 2) untreated control (n = 8). Untreated control catheters were flushed with heparinized saline daily. After completion of each daily treatment with ACL, the solution was removed and the ACL – treated catheters were flushed with 300 μl of heparinized saline. The animals were then euthanized with a pentobarbital cardiac injection and the catheters removed for quantitative culture and scanning electron microscopy (SEM) as described earlier (24).

### Quantitative catheter culture

Catheters were cut into segments. These segments were cut longitudinally and placed in sterile saline (10 ml). These segments were sonicated at 40,000 Hz (Bransonic 1510; Branson Ultrasonics Corp., Danbury, CT) for 12 min at 4-min intervals and vortexed for 15 sec. Suspensions were serially diluted and 1-ml aliquots were spread onto plates containing Sabouraud dextrose agar (Difco Laboratories) supplemented with chloramphenicol and gentamicin. Plates were incubated for 48 h at 37°C, and CFUs were counted. Catheters resulting in ≥ 100 CFUs were considered infected, based on previous studies (25).

### Scanning electron microscopy

Catheter pieces were fixed (in 2% glutaraldehyde), dehydrated, sputter-coated with gold-palladium (60/40), and viewed under a Phillips XL30 scanning electron microscope, as described in our previous publications (26, 34).

### Statistical evaluation

The mean CFU from quantitative catheter cultures were compared by the T test using SPSS version 24 software (IBM Corp, Armonk, New York). A *P* value of > 0.05 was considered significant.

## RESULTS

### Activity of ACL against catheter-associated biofilms *in vitro*

The data demonstrate that ACL was able to prevent biofilm formation by resistant and susceptible *E*. *coli*, *E*. *faecium*, *P*. *aeruginsa*, as well as MRSA and *C*. *albicans* after a 15-min exposure (CFU = 0 for each organism) (Figure 1). ACL was also able to prevent biofilm formation against resistant *-E*. *cloacae* (n=1), and *S*. *marcescens* (n=1) after 15-minute exposure (CFU = 0). VRE biofilm was the most resistant of the species tested. ACL treatment of VRE discs resulted in a 1.75 log reduction of CFU compared to PBS control discs at 15 minutes and 3.09 log CFU at 30 minutes of exposure (log CFU of 1.32 and 0 vs. 3.07 and 3.09, respectively). Increasing the exposure time of ACL to VRE biofilms to 60 min resulted in complete eradication and zero CFUs outgrowth.

**Figure 1.**
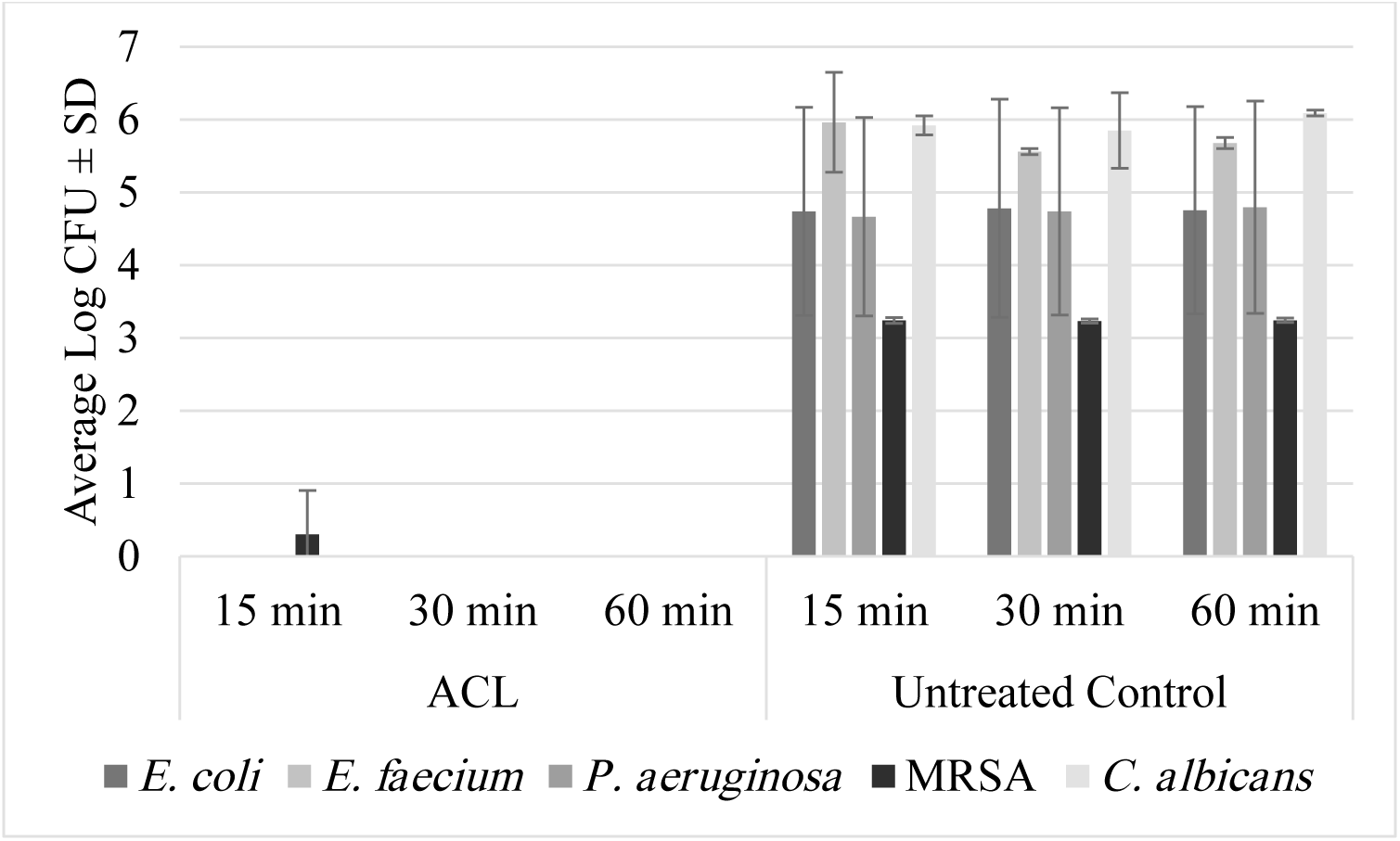
Average log CFU ± SD of ACL and untreated control disks agaisnt adhesion phase biofilms formed by trimethoprim-susceptible and trimethoprim/multi-resistant *E*. *coli* (n=3), *Enterococcus faecium* (n=2), *P*. *aeruginosa* (n=3), MRSA (n=4) and *C*. *albicans* (n=3).

Exposure of mature biofilms formed by resistant and susceptible *E*. *coli*, *E*. *faecium*, *P*. *aeruginsa*, as well as MRSA and *C*. *albicans* to ACL for 2 or 4 hours also resulted in eradication of biofilms, compared to PBS control, with ACL–treated disks resulting in 0 CFU, while the PBS control disks yielded an average log CFUs of 5.25 – 6.73 (Figure 2). Against one strain each of resistant-*E*. *cloacae*, *S*. *marcescens*, and VRE ACL was also able to eliminate mature biofilms after 2 and 4-hour exposure. The ACL–treated disks all had 0 CFU, while the PBS treated control disks yielded log CFUs of 4.43 – 6.15. Figure 3 shows representative images of the effect of ACL on adhesion and mature phase *S*. *marcescens* bacterial biofilms *in vitro*.

**Figure 2.**
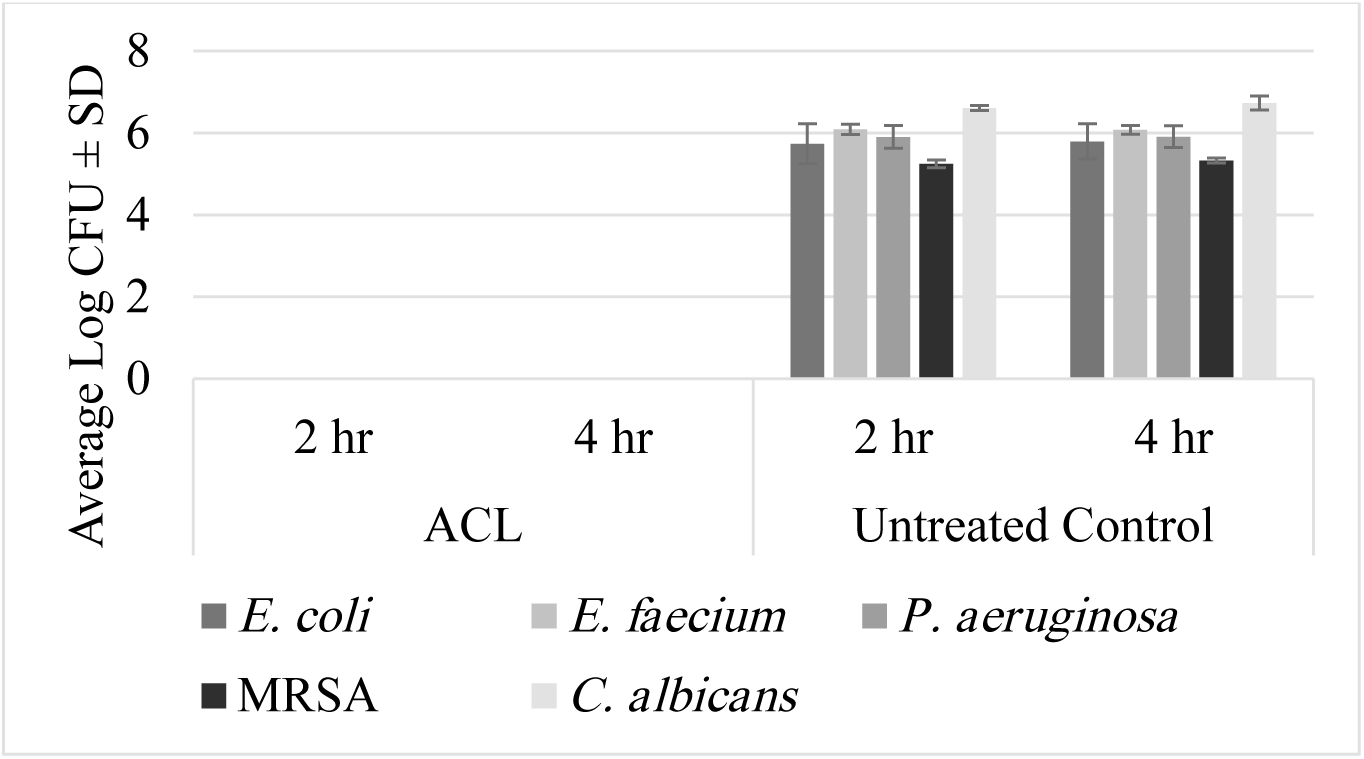
Average log CFU ± SD of ACL and untreated control disks against mature biofilms formed by *E*. *coli* (n=3), *Enterococcus faecium* (n=2), *P*. *aeruginosa* (n=3), MRSA (n=4) and *C*. *albicans* (n=3).

**Figure 3.**
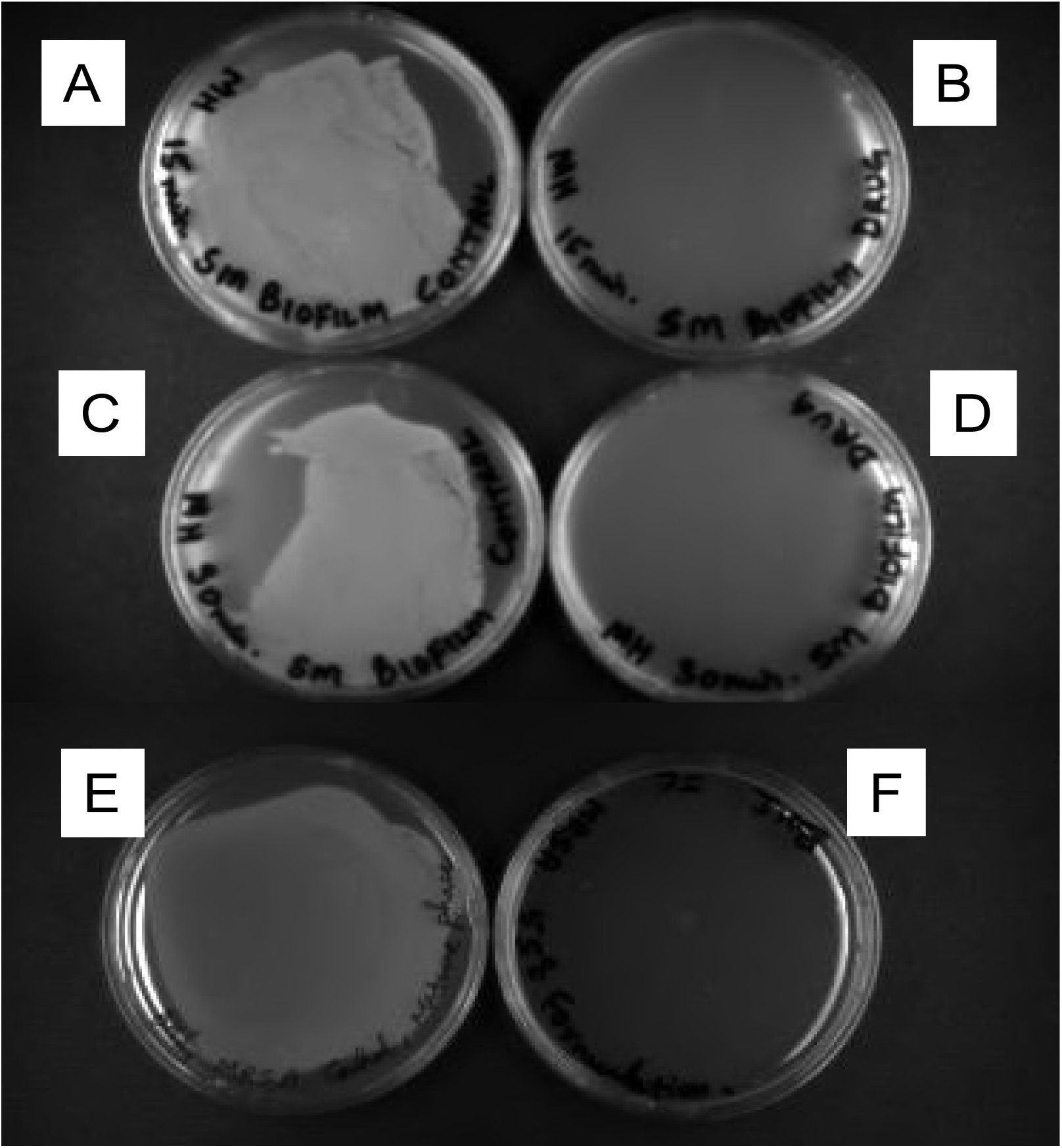
Activity of ACL against adhesion and mature phase biofilms formed by *Serratia marcescens in vitro*. (A, C, E) untreated controls; **(B, D)** biofilms exposed to ACL for 15 or 30 min, respectively; (F) mature biofilms exposed for 4h to ACL;

### Efficacy of ACL against catheter-associated biofilms *in vivo*

Next, we evaluated the *in vivo* efficacy of ACL against bacterial and fungal biofilms. For evaluation of efficacy against bacterial biofilms, blood samples obtained from the rabbit model catheters on day 3 post-inoculation (prior to the initiation of lock therapy) were positive for MRSA. This confirmed the presence of catheter-associated MRSA biofilm infections. Untreated controls yielded an average bacterial burden of 3.49 ± 0.75 log CFU/catheter (Figure 4). In contrast, ACL–treated catheters yielded an average bacterial burden of 0.03 ± 0.06 log CFU/ catheter (*P* = 0.001, Figure 4). Lock therapy with ACL completely cleared MRSA infections in three of four catheters, yielding zero CFUs in each (the fourth yielded 0.13 CFUs). SEM images showed eradication of biofilms from catheters exposed to ACL, while heavy biofilms with typical structural and architectural features were observed on untreated control catheters (Figure 5A, B).

**Figure 4.**
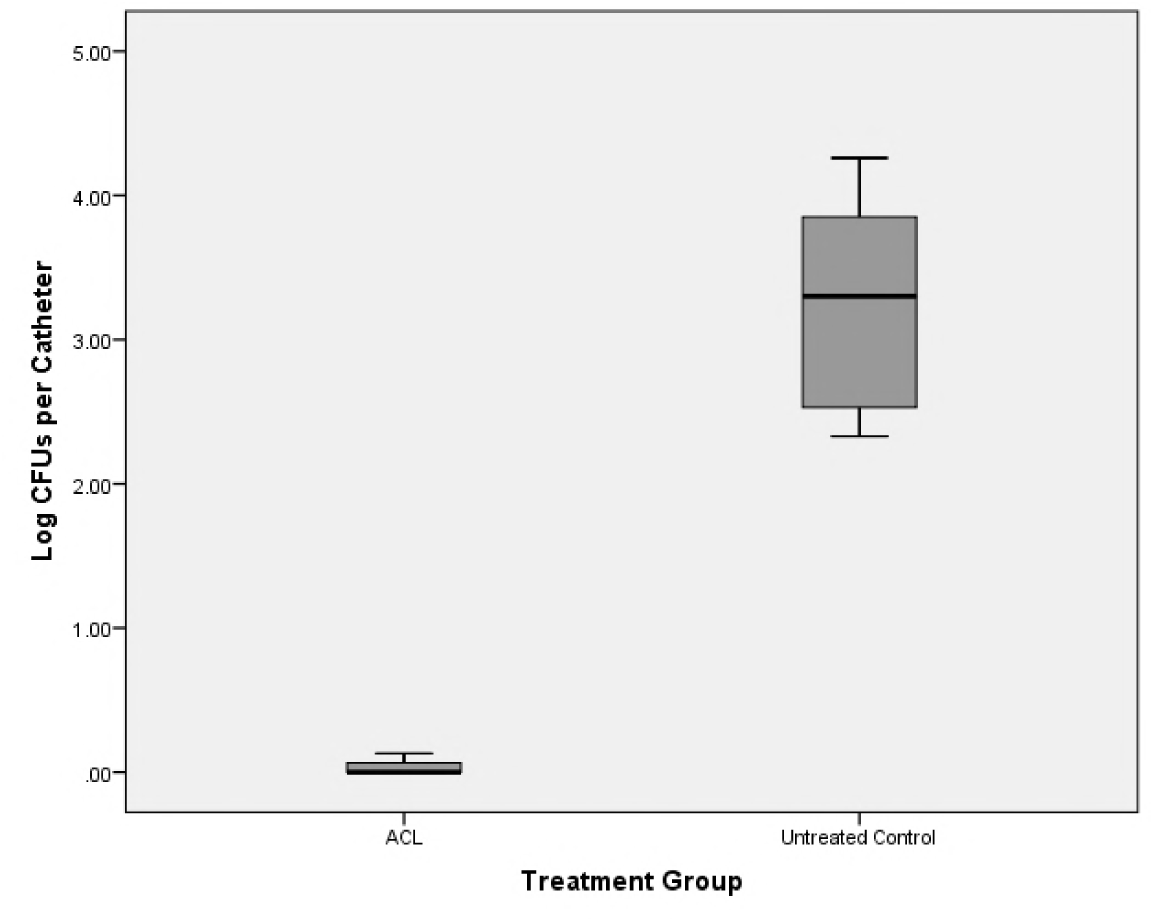
Bacterial burden of catheters obtained from animals treated with ACL and untreated control animals inoculated with methicillin-resistant *S*. *aureus* biofilms *in vivo*.

**Figure 5.**
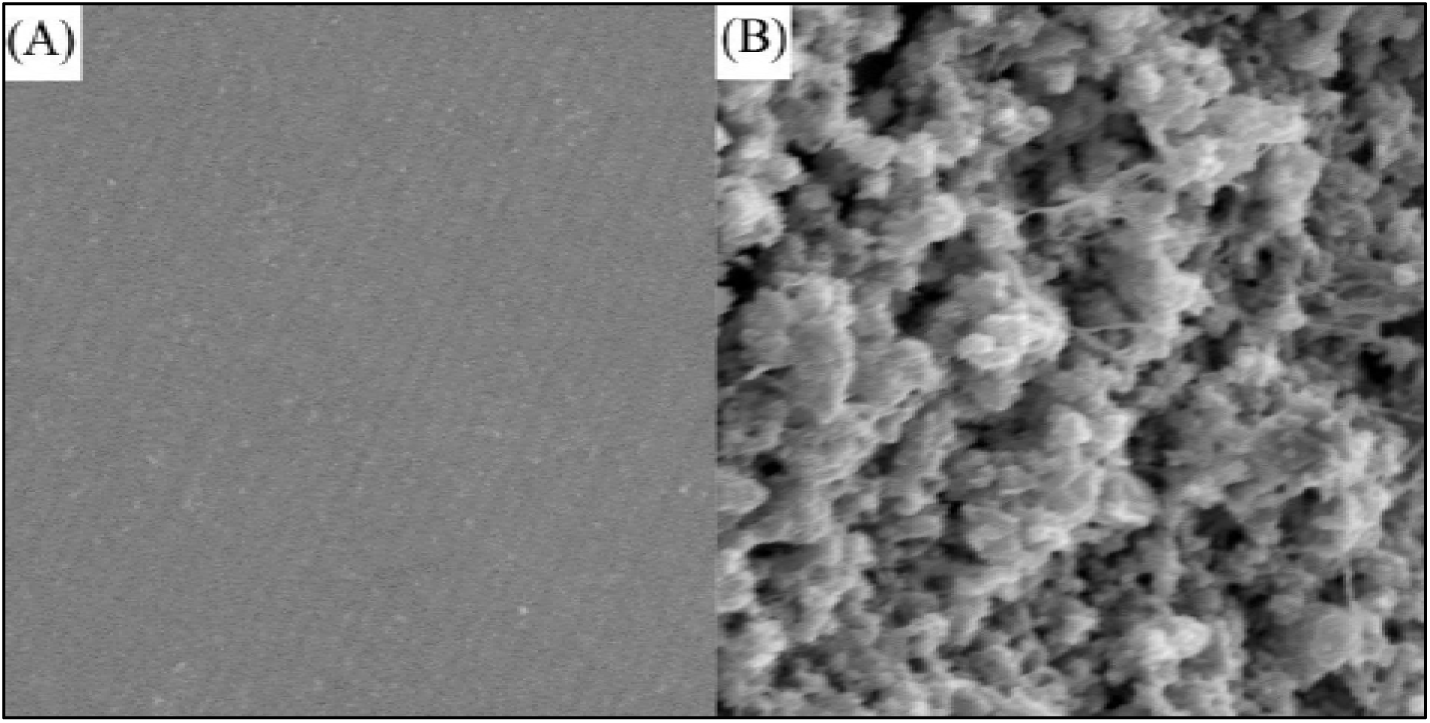
Efficacy of ACL against methicillin-resistant *S*. *aureus* biofilms *in vivo*. Representative scanning electron micrographs of intraluminal surface of catheters obtained from (A) animals treated with ACL or (B) untreated animals.

For evaluation of antifungal biofilm efficacy, blood culture samples obtained from the catheters on day 3 post-infection (prior to the initiation of lock therapy) were positive for *C*. *albicans*, again confirming the presence of catheter associated biofilm infections. There was a significant difference in the mean log CFUs between animals treated with ACL (0.55 ± 1.19 log CFU/catheter segment) and untreated control (2.48 ± 1.60 log CFU/ catheter segment, *P* = 0.012) (Figure 6). Lock therapy with ACL completely cleared fungal cells in eight catheters (zero CFUs in each).

**Figure 6.**
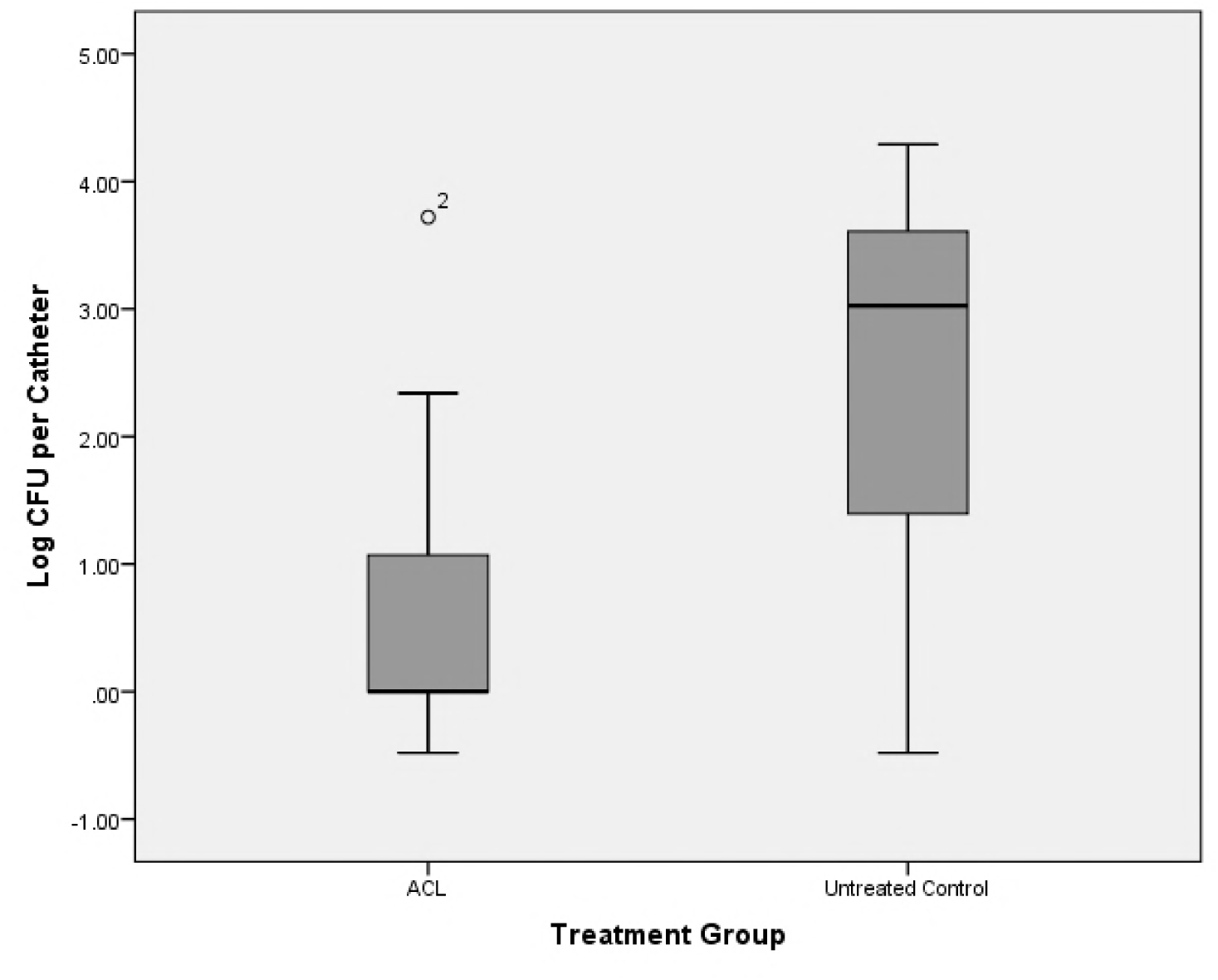
Fungal burden of catheters obtained from animals treated with ACL and untreated control animals inoculated with *C*. *albicans* biofilms *in vivo*.

## DISCUSSION

These studies demonstrate that ACL successfully prevented formation of biofilms and also eradicated mature biofilms formed by multidrug resistant bacteria and fungal species known to cause catheter blood stream infections in humans. Previous studies have reported variable antimicrobial activity of ethanol and EDTA, alone or in combination. In this regard, Passerini de Rossi *et al*. (27) showed that exposure of mature biofilms formed on silicone catheter to ethanol (25 or 40%) or a combination of ethanol (25%) and EDTA (30 mg/ml) for 1 h resulted in inhibition of biofilm growth. However, complete eradication was not observed for EDTA treatment, even after exposure for 24 h (27). In a separate study, Parra *et al*. (28), reported that antibiotics (daptomycin, teicoplanin, clarythromycin) combined with ethanol (35%) significantly reduced biofilms formed by methicillin resistant coagulase negative staphylococci after 72 h exposure. These studies are in agreement with our finding that combination treatment with an antibiotic and ethanol in an aqueous solution is effective against mature biofilms formed by Gram-positive, Gram-negative bacteria and fungal pathogens.

Although there is increasing recognition that antibiotic lock solutions can reduce the risk of infection, the standard lock solutions used clinically remain PBS alone or containing Heparin as an anti-coagulant agent Chapla *et al*. (29). Heparin is a sulfated polysaccharide analogous to carrageenan, also a sulfated polysaccharide that is isolated from marine algae and used as a growth substrate in microbiology plates to culture bacteria (10). Moore *et al*. (30) compared the efficacy of gentamicin/citrate and heparin catheter lock solutions and reported significant reduction (73%) in the rate of BSIs and mortality. A limitation of using off-label gentamicin / citrate lock solutions is that rather than being microbicidal it is only bacteriostatic and resulted in increased recovery of gentamycin-resistant bacteria in the treatment cohort compared to the heparin arm. Murray *et al*. (31) reported that introduction of taurolidine-citrate-heparin resulted in a significant reduction (56%) in staphylococcal bacteremia in haemodialysis patients. Saxena *et al*. (32) showed that cefotaxime-heparin locks for tunneled cuffed catheters reduced the incidence of catheter related BSIs associated with methicillin-susceptible *S*. *aureus* (MSSA) but did not show activity against MRSA. Overall, these investigations have led to the suggestion that heparin lock solution should be abandoned due to the risks of rapid biofilm growth and bleeding (33).

Results from this study expand on previous publications by demonstrating that ACL was effective in preventing adhesion phase cells from developing into mature biofilms and also effectively at eradicating fully formed mature biofilms. Our results also show that ACL was able to eradicate bacterial and fungal biofilms formed *in vivo* after seven days of lock therapy. This novel lock solution may provide important advantages in preventing the introduction of skin flora or other nosocomial infections into the bloodstream of patients with indwelling catheters that must be routinely handled by health care professionals. Further, use of this novel lock solution may help avoid the risk and added expense of removing infected indwelling catheters. Based on these findings, further clinical testing of this novel lock therapy solution is warranted.

## ACKNOWLEDGMENTS

Funding support is acknowledged from NIH (R01DE024228) to MAG and PKM, and the Steris Foundation (Research Award to PKM).

